# Measuring anesthetic resistance in Drosophila by VAAPR

**DOI:** 10.1101/797209

**Authors:** E. Nicholas Petersen, Katherine R. Clowes, Scott B. Hansen

## Abstract

Volatile anesthetics are compounds which are commonly used to induce a reversable loss of consciousness (LOC) in animals. The molecular mechanism of how anesthetics induce LOC is largely unknown. However, observations have been made which show that there are genetically-encoded traits which influence the effective concentration of anesthetics in the inducement of LOC. Despite this long-term observation, little progress has been made in identifying genes involved in anesthetic sensitivity. One reason for this is that many techniques to test anesthetic sensitivity are technically challenging and are inhibitory for high-throughput studies. Here we introduce a technique for testing volatiles and aerosols with positional recording (VAAPR), a method which allows for high-throughput testing of the effect of anesthetics and other aerosolized drugs using Drosophila. Using VAAPR we show that the enzyme phospholipase D (PLD) significantly shifts the concentration of diethyl ether, chloroform, and isoflurane needed to induce LOC in Drosophila. We also show that PLD is required for a paradoxical hyperactivity phenotype. We expect that this technique will allow for additional genes to be found which control anesthetic sensitivity as well as other behavioral phenotypes.

## Introduction

Since the early 19^th^ century volatile anesthetics have been used to cause temporary loss of consciousness in an individual, normally so that invasive medical procedures could be performed^1^. Ether was the first volatile anesthetic used during surgical purposes, followed quickly by chloroform, and later by the flurane-derived family of anesthetics. Since their discovery, the mechanisms of action have been of great interest, largely due to the fact that such a diverse array of compounds could elicit the same phenotype throughout multiple animal kingdoms.

Recently, experiments have shown that volatile anesthetics can have a direct effect on the secondary organization of the lipids in the plasma membrane^2–4^. The enzyme phospholipase D (PLD) is known to be regulated by localization to lipid domains^5^, and disruption of these domains by anesthetics have shown activation of the enzyme^2^. Since PLD is also known to bind to and regulate the activity of associated ion channels through the production of the signaling lipid phosphatidic acid (PA)^6,7^, we wanted to know if the activation of PLD by anesthetics could play a role in the temporary loss of consciousness commonly attributed to volatile anesthetics. We chose to answer this question using D. melanogaster since this animal only has one copy of the PLD gene and exhibits similar effects to anesthetics as their mammalian counterparts^8^.

In order to best answer this question, we developed a technique that would allow for fast, high-throughput, scalable, and repeatable observation of the phenotypes associated with anesthetic application. In order to achieve this, we made a video tracking -based assay as this would allow for multiple variables to be assessed while the drug was being administered. Other techniques have been used to measure sedation in flies but many of them require large cohorts of flies or use limited observation technologies (e.g. beam breaks, death under sedation) limiting the data obtained^9–13^. The high Hill slopes of many of these anesthetics also make it difficult to discern minute changes in sensitivity which may exist between two genotypes of flies requiring high sensitivity to observe changes.

Using a new video-tracking technique we call volatiles and aerosols with positional recording (VAAPR) we can directly measure the effect of volatile anesthetics on the sedation of flies. We show that with this technique we can measure the anesthetic potency of chloroform, isoflurane, and diethyl ether on flies. We also show that flies without functional PLD (PLD^null^) were resistant to both the sedative and the paradoxical hyperactivity state of all these anesthetics.

## Results

### Protocol for anesthetic application to flies

To test the effect of anionic lipid on the effects of anesthetics we developed the volatiles and aerosols administers with positional recording (VAAPR) to accurately measure the physiological response to varying dosages of drugs of interest on fruit flies. VAAPR consists of a 3D printed device with narrow, vertical-width columns covered with a clear acrylic sheet in order to monitor the activity of the flies contained in each lane with a webcam. A single fly is placed into each column using an aspirator and flies are allowed to acclimate to the chamber for at least 5 min. Once acclimation is complete, flowmeters are used to control the air or air/anesthetic flow into the chambers (Fig. 1a). The flow meters allow both for control of total air volume as well as the dosage of anesthetic being applied.

**Fig 1.**
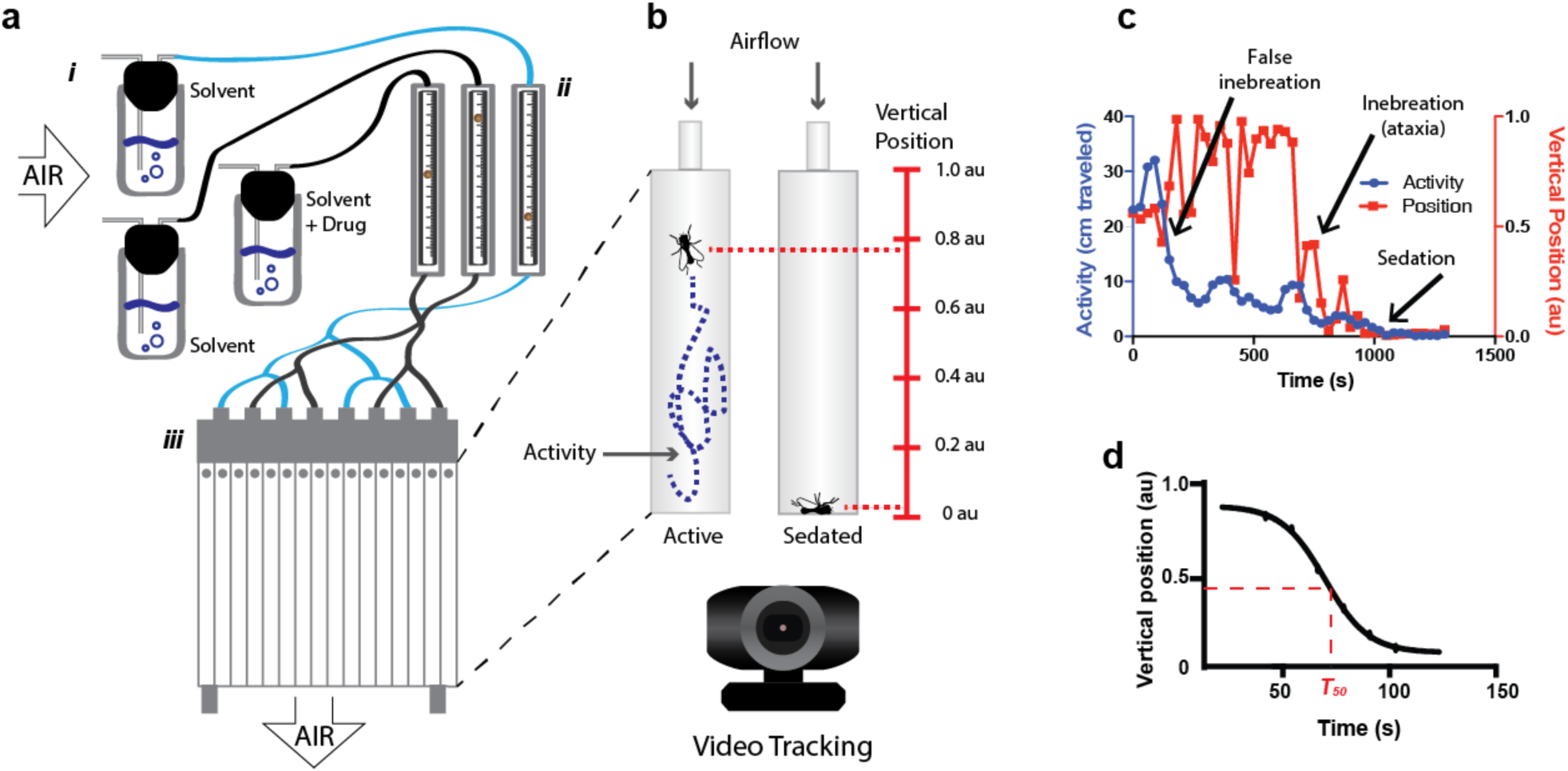
High throughput method for measuring the effects of volatile drugs. (a) Example setup for VAAPR. Air is pushed through vials containing a solvent and/or an aerosolizable or volatile compound(i). The vapors are passed though flow meters which control the total concentration of drug administered (ii). By using more than one flow meter the total flow rate of air can be kept constant while the concentration of compound is varied. Finally, the treated air is passed into a imaging chamber (iii) where effects on individual flies can be measured. (b) The behavior of flies can be monitored using a webcam which allows for measuring of the instantaneous activity level (left, blue) and position (right, red) for each fly to be measured. Analysis of the data for an individual fly (c) illustrates the ability to use the activity and position data together to gather information on the overall state of the fly at any one specific time. To get a measure of sedation, a T50 (time to half position) is measured by fitting a curve to the position graph of an individual fly (d). This value can be combined with other flies to obtain an average time to sedation for a specific concentration of compound.

Individual flies are tracked during drug application for both activity and position using a webcam and custom software developed by the Ja lab^14^. Addition of a mildly aversive odor (methylcyclohexanol, MCH) to the bulk air flow allows for a baseline of activity and a randomized position to be retained during airflow (Supplemental Fig. 1). For each run flies were loaded into the chamber with experiments containing more than one genotype having each line put into alternating lanes to decrease variability due to slight differences in maze position/air flow. For each experiment flies were subjected to 2 min of baseline air (air + MCH) after which the anesthetic of interest was dosed into the chambers. This was allowed to continue until all the flies were observed to have lost consciousness after which the tracking was stopped and the flies discarded. Depending on the dose of anesthetic given each run would last from 5 min to an hour.

Tracking both variables of position and activity allows for more robust determination (less false positives) for the time of sedation. Anesthesia was found to result in a steep, persistent decrease in the position value of each animal to zero whereas activity would decrease slowly until reaching a minimum level with or without sedation. For this reason, position was used as the main marker of LOC with activity allowing for secondary confirmation of LOC. Using curve-fitting software, a time of sedation can be determined from the position graph for each fly empirically resulting in an individual sedation time for each animal (T50 value, Figure 1d). Each T50 value was then put into a survival curve to allow for statistical determination of LOC time as well as allowing for comparison of sedation between experimental groups. After determining that this method of anesthetic application and data analysis could robustly determine the time of LOC we then desired to determine if VAAPR could be used to measure changes in anesthetic sensitivity between experimental groups.

### Effect of PLD on the sensitivity of volatile anesthetics

We found that loss of the protein PLD resulted in a resistance to anesthesia. Background-matched WT flies (w^1118^) were compared to flies without a functional PLD (PLD^null^). We then applied increasing concentrations of anesthesia until the time to LOC had reached a minimum. Concentrations were measured as the concentration applied and not the concentration which existed in the flies at the time of sedation. We performed this comparison between w1118 and PLDnull flies using chloroform, diethyl ether, and isoflurane. (Figure 2).

**Fig 2.**
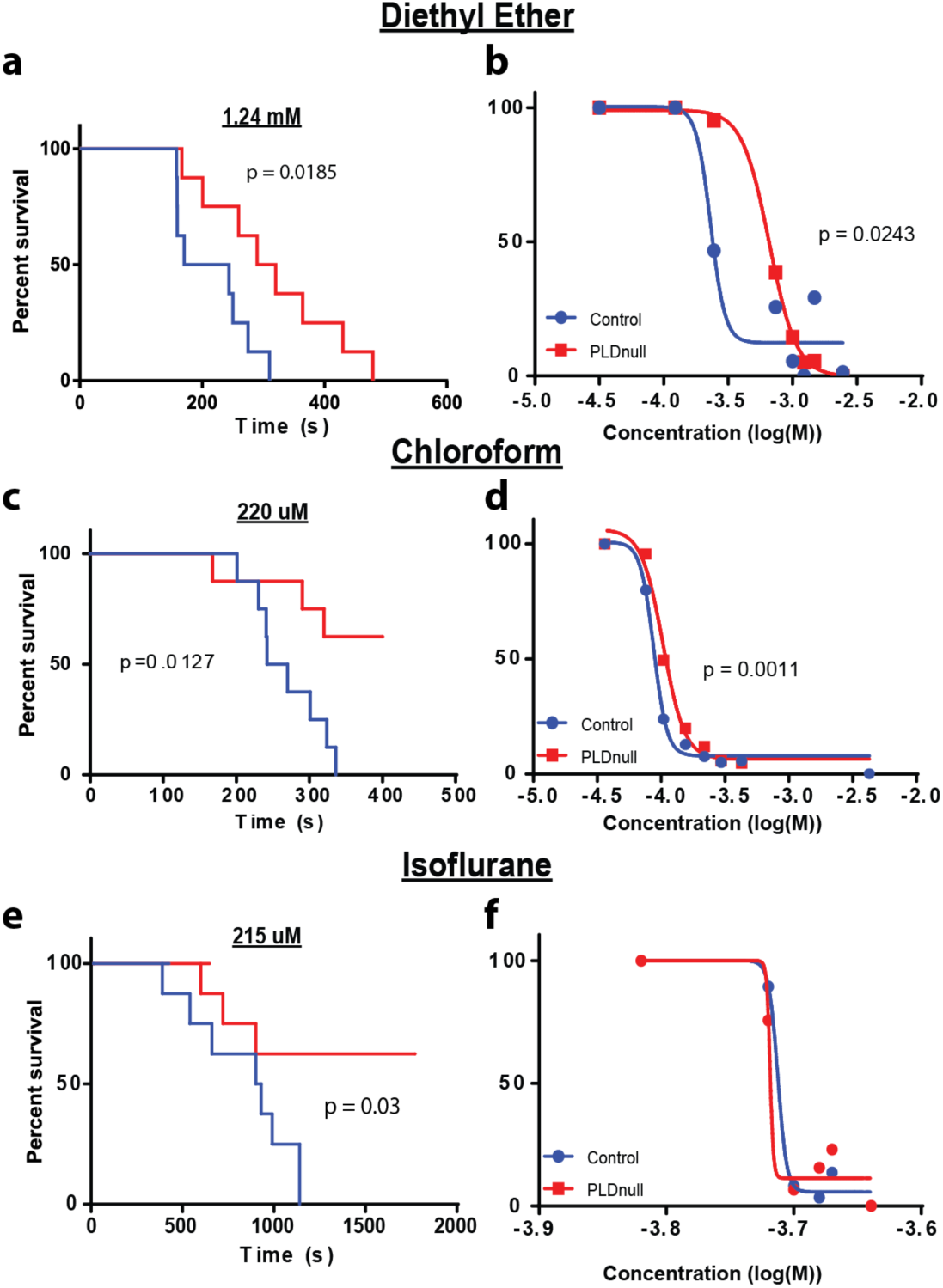
Effect of PLD^null^ mutant on anesthetic sensitivity. Single consciousness curves (left) and full dose response curves (right) for diethyl either, chloroform, and isoflurane. Consciousness curves for diethyl either(a) chloroform (c) and isoflurane (e) all show significant changes in the time it takes the population of flies to become fully sedated. Comparing the EC50 between the two groups, diethyl ether was very sensitive to the loss of PLD functionality in the fly resulting in a significant rightward shift of the curve (b). A significant shift was also noted for chloroform (c), although the magnitude of the shift was greatly decreased. Isoflurane did not show a significant shift in the overall determination of EC50 (f). Each point on these graphs was determined with the n = 8-16 over a minimum of two runs. P values are noted where significant.

In all cases we observed a resistance to anesthesia in the PLD^null^ flies as we would have predicted. The ED50 of diethyl ether shifted the most with a 2.8-fold shift from 230 µM to 660 uM (Fig. 2a). The change observed for animals in chloroform was 1.2-fold from 88 µM to 104 µM (Fig. 2b). The shift in isoflurane was not significant when calculating the overall ED50 (Fig. 2c), most likely due the extreme Hill slope (∼200) making it difficult to obtain much precision. Despite this we were able to find multiple single concentrations at which PLD^null^ flies showed resistance to isoflurane (Supp. Fig 4), matching the effects seen in chloroform and diethyl ether.

### Using VAAPR to study the hyperactive effect of anesthetics

Many anesthetics including alcohols, propofol, and volatile anesthetics have been shown to cause hyperactivity in animals preceding their sedative effect^15–18^. While the hyperactive effect during alcohol consumption has been shown to rely on PLD^18^, little is known about the cause of hyperactivity, or ‘paradoxical state’ for the other anesthetic families. It has been proposed that the pre-anesthetizing pain felt during local anesthetic application (e.g. lidocaine) may be dependent on their interaction with PLD and an associated ion channel^19^. We monitored the hyperactive state during volatile anesthetic application using VAAPR and determine if this effect was PLD-dependent.

We found that w1118 flies did show a hyperactive response preceding anesthesia. We wrote a custom program which allowed us to normalize the activity tracking of each fly to the moment of LOC. Then we fit a 4^th^ order polynomial trendline to the data to allow for unbiased visualization of any hyperactive peak working opposite the normal activity decline. For all anesthetics, an increase in activity was noted before LOC occurred (Fig. 3, blue traces with arrow). This hyperactivity was only found in the concentration for which each respective anesthetic was at a minimum but still caused all flies to be sedated. At higher concentrations this effect is likely masked due to the quick onset of LOC.

**Fig. 3.**
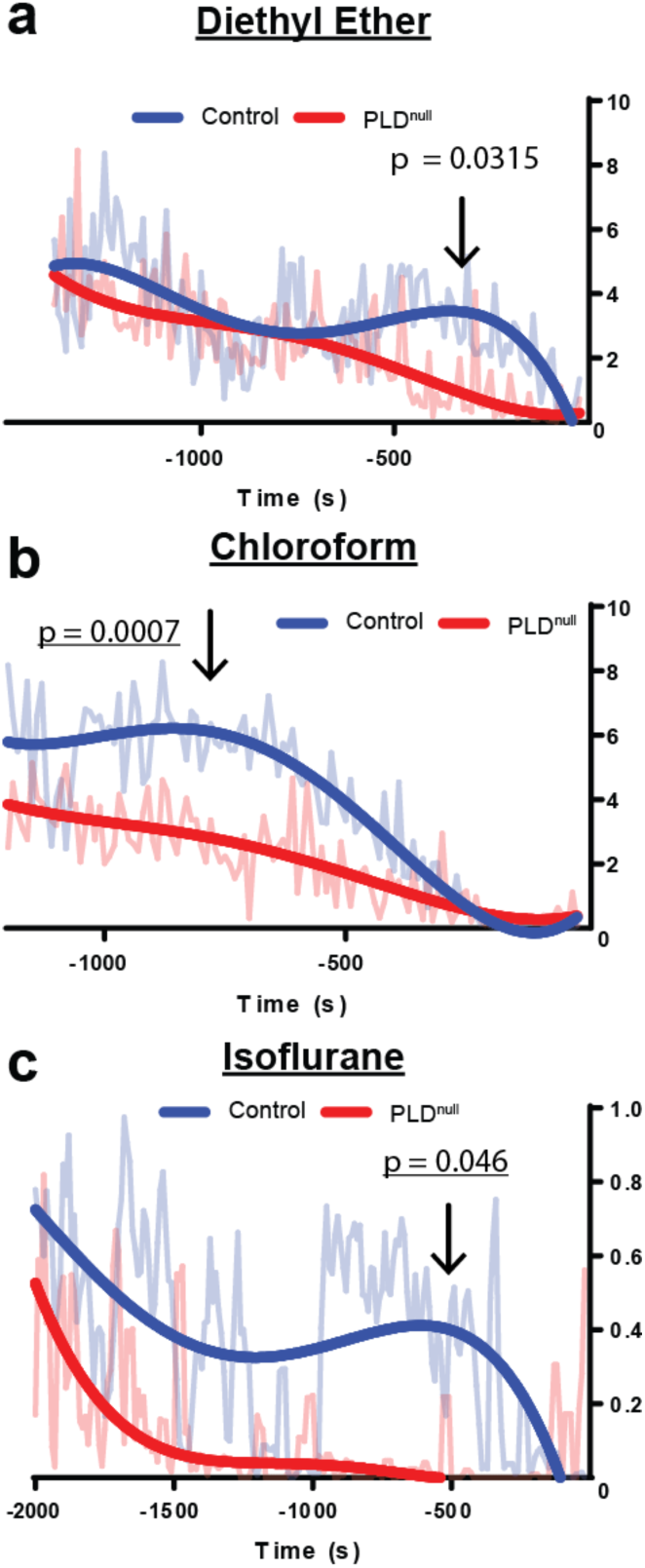
PLD^null^ mutant abolishes the hyperactivity phenotype of volatile anesthetics. Activity traces with the time 0 corresponding with the onset of LOC for all flies. (a-c) For WT flies, a small increase in activity (arrow) was noted preceding LOC but after application of anesthetic (time not shown). This effect was not noted in PLD^null^ flies. All traces are an average of activity for 8-16 flies, 10 sec bins. Trendline is best fit to the average with a 4^th^ order polynomial. P values are calculated by comparing the total activity between control and PLD^null^ flies under the hyperactivity curve (n = 7-8).

PLDnull flies did not have a hyperactivity peak before LOC. The data for these flies was normalized and fit with the same trendline, but no significant hyperactivity peak was observed for any of the anesthetics (Fig. 3, red traces).

## Discussion

We have shown here that VAAPR is a viable, high-throughput, high-resolution technique for analyzing the behavior of flies in response to an aerosolized or volatile drug. While the chamber built here holds 16 flies, it would be easy to scale this up to achieve much greater throughput. The combination of video tracking along with control over the airflow with digital flowmeters allowed for extremely precise and repeatable measurements to be made. While this is needed to study the EC50 of drugs like anesthetics, analog flowmeters could be used to study compounds whose effect does not have as steep of Hill slopes or whose magnitude of observed phenotype is larger.

VAAPR has allowed us to determine that production of anionic lipid by PLD is important in controlling both the threshold of the loss of consciousness and the preceding hyperactive state during anesthetic application. PLD is known to be modulated by anesthesia^19^. Rather than completely inhibiting or leading to spontaneous activation of loss of consciousness, the production of PA seems to be important in controlling the threshold at which this event occurs. It is unclear at this point whether this threshold control is happening at a single-cell level or whether this is a threshold at the systems level, but more specific control of PLD knockdown may be used to help with this determination. It is unknown how PLD elicits the hyperactive response, but it is likely by effecting downstream, lipid-sensitive excitatory ion channels whose Kd for PA is slightly lower than the inhibitory channels PLD is known to affect5.

PLD is known to be sensitive to its localization to lipid domains^5^ and recent research has shown that anesthetics have a significant effect on the organization of these domains^2,19,20^. PLD and its activating lipid PA have also been shown to directly control the activity of the mammalian potassium channel TREK-1, a hyperpolarizing channel^7,21^. While it is unknown what channels with which the fly ortholog of PLD are associated, recent experiments have shown that it too is likely helping to regulate a hyperpolarizing channel like TREK-1.

By controlling the threshold of activation, PLD offers an interesting method for possibly modulating the activation of the pathway controlling loss of consciousness. Some sleep disorders such as insomnia may have roots in a lack of ability to initiate the sleep cycle, and while anesthesia-induced sleep does not recapitulate all the aspects of natural sleep^16^, there is commonality in that both begin with an initial loss of consciousness. It may be that this pathway is shared and that anesthetics offer an alternative method by which to study this important pathway.

## Acknowledgements

We thank William Ja, Keith Murphy, Scarlet J Park (Scripps Research) for help with software development and helpful conversation. This work was supported by a Director’s New Innovator Award (1DP2NS087943-01) and an RO1 (1R01NS112534-01) from the National Institutes of Health.

## Materials and Methods

### Materials

#### VAAPR

PLA, VWR 470221-854; webcam, Microsoft LifeCam Studio; y-connectors, FisherSci 50630702; viton tubing, Cole-Parmer 06434-04; PTFE tubing, Grainger 2VLW9; flowmeters, Alicat Scientific MCS-5SLPM & MC-5SLPM; acrylic sheet (0.93 inch), Optix.

#### Anesthetics

Isoflurane USP, Patterson Veterinary; Chloroform, FisherSci C606-4;Diethyl either, Acros organics 61508-5000

### VAAPR chamber

The VAAPR system consists of 2 major parts, 1) the flowmeters and 2) the treatment chamber.

1. To set up the flowmeter section, air is forced through bottles one of which contains the air/MCH mixture and the other which contains anesthetic/drug. In our setup we used a 250 mL bottle with a rubber cork inset with two custom-bent glass tubing for the air/MCH and a 25 mL bottle with the same setup for the anesthetic/drug. These can be changed to suit the needs of the experiment. The air from these bottles was then pumped into flowmeters where the overall flow rate was adjusted to apply the concentration of drug as desired. Total flow rate was kept constant and the concentration of drug given was calculated using the partial pressure of each anesthetic. Due to the fast-evaporation of the anesthetic leading to cooling over time, anesthetic vial was kept in a large volume of water to maintain the desired temperature. From the flowmeters the air was combined into a single tube using a Y-connector which was then extended to the chamber where it was then split into 8 outputs using Y-connectors. The splitter was connected to the 8 outputs from the flowmeter.
2. A printed chamber with 16 total lanes was printed and topped with a splitter to allow for 8 inputs to split into 16 individual chambers. Between the chamber and the splitter at at the bottom of the chamber was attached (using chloroform) with fine wire mesh to block egress of the flies. The chamber was held using custom-printed legs and U brackets. The U brackets were glued to a piece of wood or other holder to allow for consistent placement of the chamber in relation to the webcam which was also held in place using a custom holder and hot glue. After placement, determination of the pixels/cm was performed so that accurate measurements of fly movement could be made.

### Drosophila

#### General protocols

Flies were maintained in stocks at 25C. For all experiments, male flies were isolated from stocks 1-4 days old and allowed to recover in vials of no more than 10 flies for 24-72 hours before use in protocols.

#### Anesthesia in Drosophila melanogaster

Anesthesia in flies were performed by volatiles and aerosols administered with positional recording (VAAPR) in a custom-designed, 3D printed device with narrow-width vertical chambers placed in front of a camera to monitor the fly’s positions with or without the chloroform treatment. Wildtype and PLD^null^ flies were gently loaded into the designated chamber using mouth aspiration and the hoses used for compound delivery were attached to the chamber. Flies were allowed to habituate in the chamber while preparing treatment mixtures (5-15 min). After mixtures are prepared, the hoses are attached to the chamber and the air/MCH mixture is started. Immediately after the video tracking is also started. Flow rates are kept constant at 144 ml/min/chamber. Air was given to all flies for 2 min to record baseline activity/position after which the anesthetic dose is applied in mixture with the air. Assay was stopped after all flies became sedated or 45 min had passed, whichever came first.

#### Data Acquisition & Analysis

Flies were tracked using a custom written program in Python, ‘opencvArduinoARC_2.0’. Data was analyzed by the ‘Noah’ program for position and activity. Both programs can be found in the ‘flyARC’ repository on Github.

### Statistics

For determination of time of sedation, we performed a non-linear fit with variable slope to determine the T50 for sedation. This value was then used for each fly to create the sedation curves. Individual curves were determined to be significant using the Mantel-Cox test. These values were then averaged to give a single point for the dose-response curves. Dose response curves were fitted using the same non-linear variable fit as above and tested for significant changes in the LogIC50 between the two lines. Hyperactivity was compared by averaging the activity underneath the curve and comparing the two averages for each phenotype with a t test. All p values are placed next to the curves in which there is significance (p < 0.05); if there is no reported p value than the p value is > 0.05. Statistics were calculated using GraphPad Prism 6.01.

---

**Supp Fig. 1.**
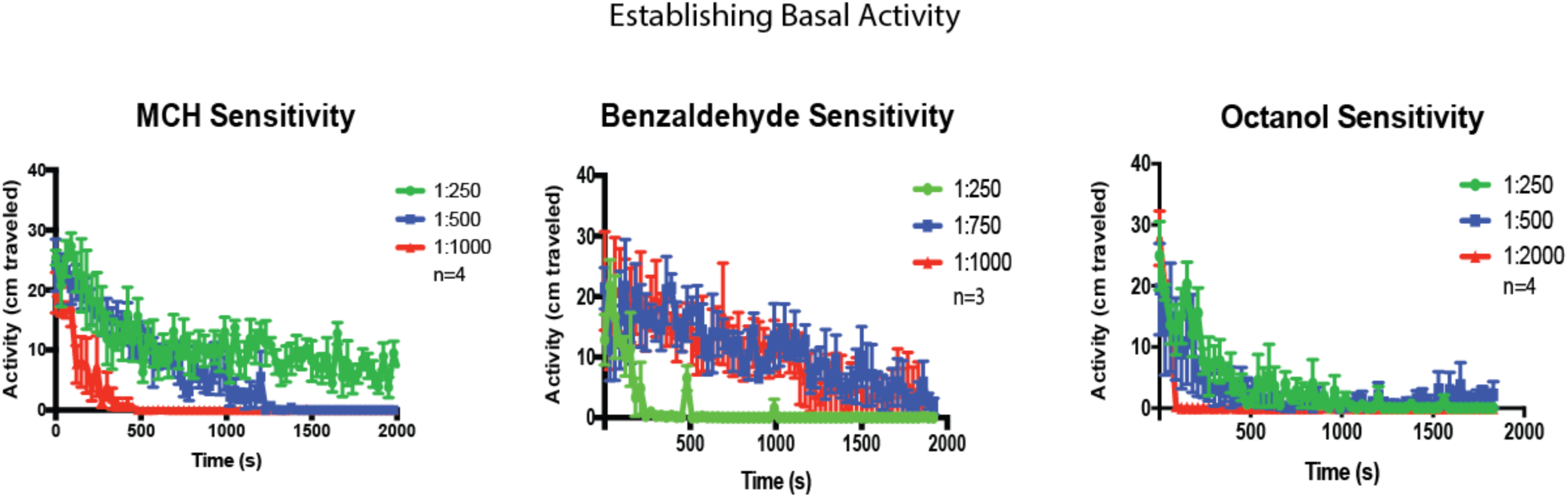
Comparison of stimulating aversive odors on individual Drosophila. 1:250 of MCH was the only aversive odor and concentration to allow for consistent activity over 30+ minutes without leading to anesthesia. At lower doses benzaldehyde and octanol were not able to maintain the activity of the flies and at higher does it seems as if they act as anesthetics since flies stopped moving but were able to recover after cessation of the odor.

**Supp Fig. 3.**
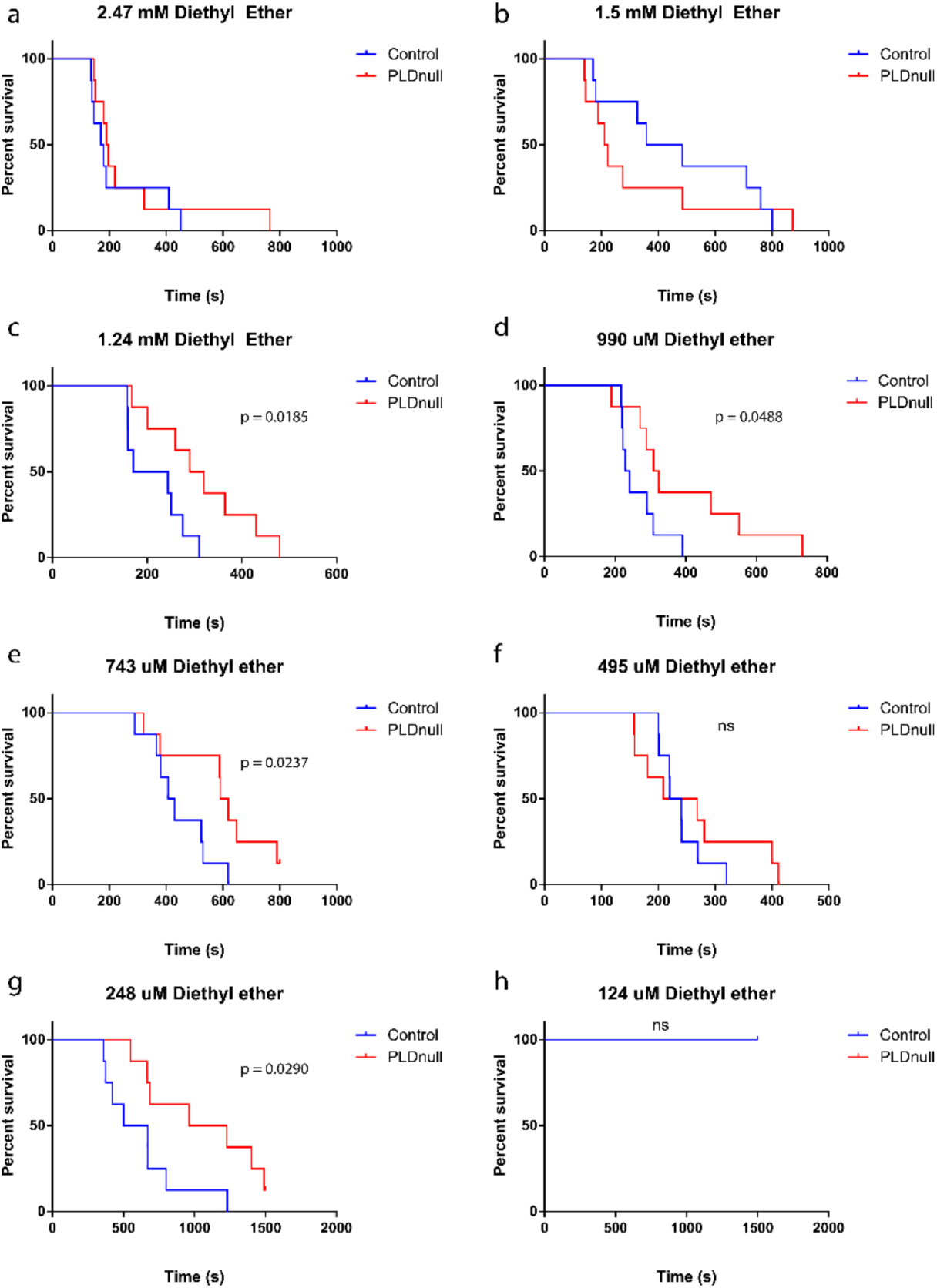
Raw survival curves for diethyl ether anesthesia. Each curve was created by taking the time each fly become sedated through observation of their overall position.

**Supp Fig. 3.**
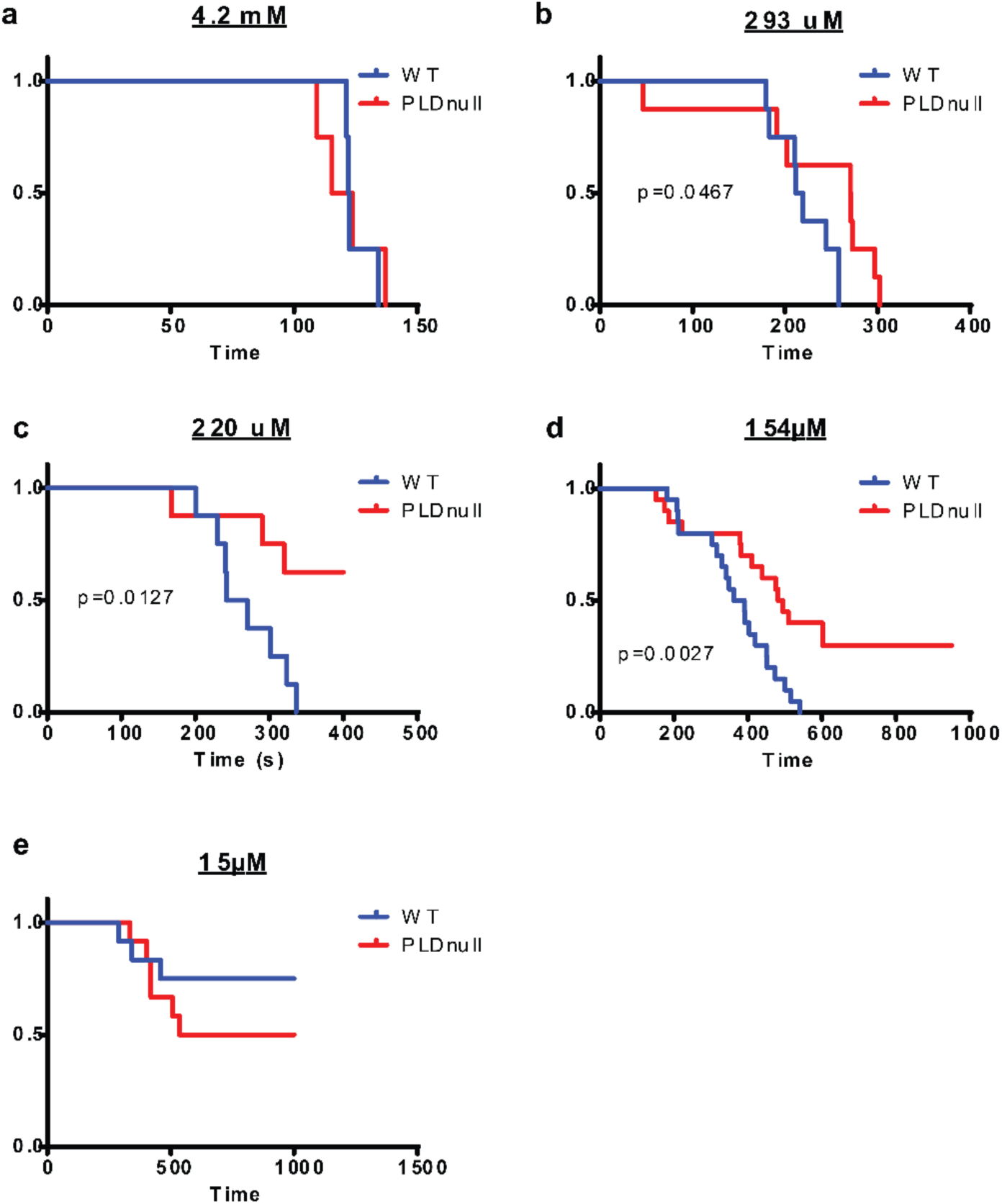
Raw survival curves for chloroform anesthesia. Increasing doses of chloroform were administered to flies and the T50 for each fly was measured and put into a sedation plot (a-e). While no differential effect between PLD^null^ and control flies was seen at very high (4.3 mM, a) or low (15um, e) concentrations, a significant effect was measured at doses in between (b-d). When plotted on a dose-response curve (f), these changes led to a significant rightward shift in PLD^null^ flies when compared to genetically matched controls.

**Supp Fig 4.**
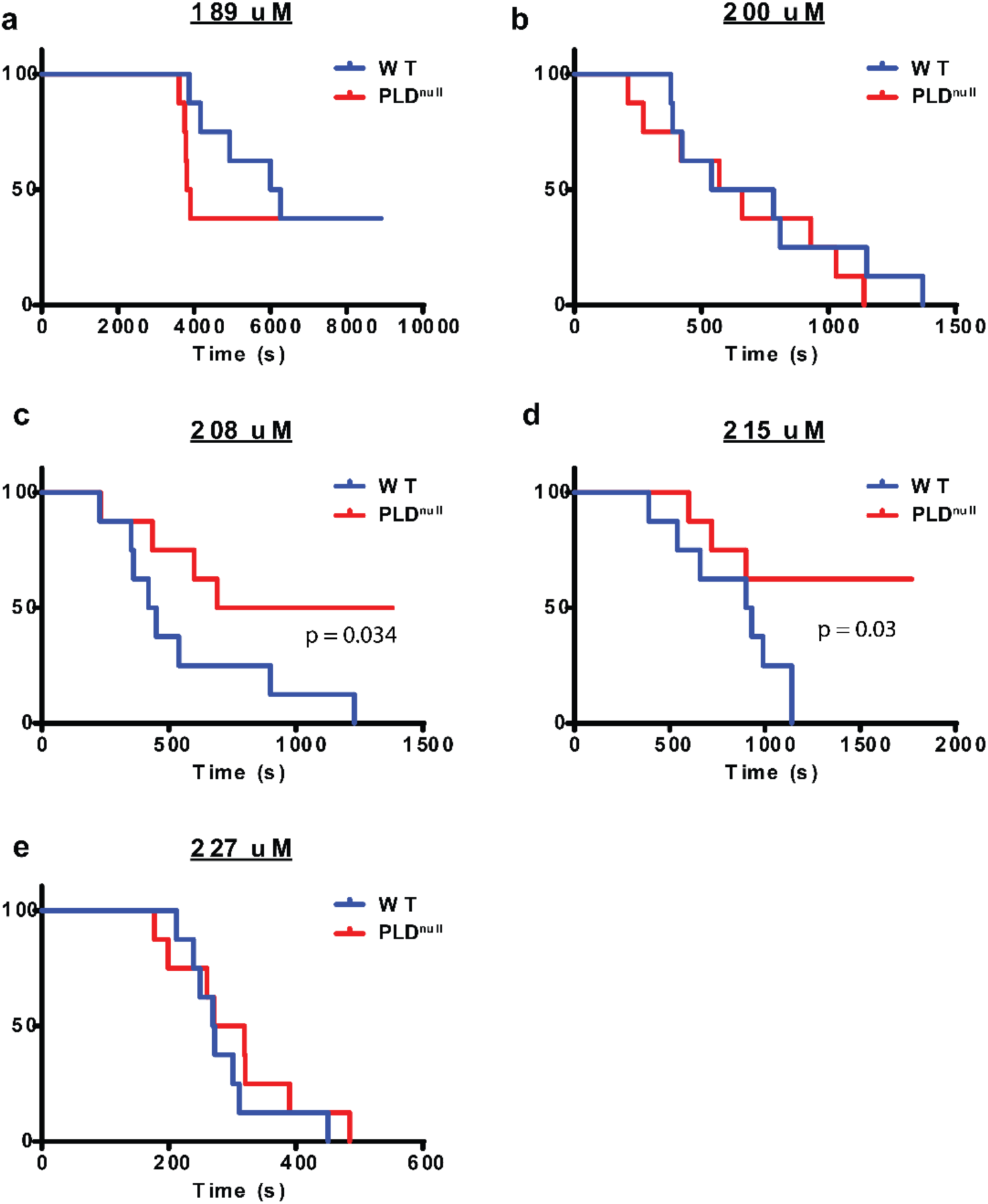
Raw survival curves for isoflurane anesthesia. Increasing doses of isoflurane were administered to flies and the T50 for each fly was measured and put into a sedation plot (a-e). Individual concentrations were shown to have significant shifts between wildtype and PLDnull flies (c-d).

